# Vaccinia Virus Arrests and Shifts the Cell Cycle

**DOI:** 10.1101/2021.07.19.452625

**Authors:** Caroline K. Martin, Jerzy Samolej, Annabel T. Olson, Cosetta Bertoli, Matthew S. Wiebe, Robertus A. M. de Bruin, Jason Mercer

**Affiliations:** MRC Laboratory for Molecular Cell Biology, University College London, London, UK; School of Veterinary and Biomedical Sciences, University of Nebraska, Lincoln, USA; Institute of Microbiology and Infection, University of Birmingham, Birmingham, United Kingdom

## Abstract

Modulation of the host cell cycle is a common strategy used by viruses to create a pro-replicative environment. To facilitate viral genome replication, vaccinia virus (VACV) has been reported to alter cell cycle regulation and trigger the host cell DNA damage response. However, the cellular factors and viral effectors that mediate these changes remain unknown. Here, we set out to investigate the effect of VACV infection on cell proliferation and host cell cycle progression. Using a subset of VACV mutants we characterize the stage of infection required for inhibition of cell proliferation and define the viral effectors required to dysregulate the host cell cycle. Consistent with previous studies, we show that VACV inhibits, and subsequently shifts the host cell cycle. We demonstrate that these two phenomena are independent of one another, with viral early genes being responsible for cell cycle inhibition, and post-replicative viral gene(s) responsible for the cell cycle shift. Extending previous findings, we show that the viral kinase F10 is required to activate the DNA damage checkpoint and that the viral B1/B12 (pseudo) kinases mediate degradation of checkpoint effectors p53 and p21 during infection. We conclude that VACV modulates host cell proliferation and host cell cycle progression through temporal expression of multiple VACV effector proteins.

## INTRODUCTION

The cell cycle is the most fundamental molecular process of all life forms, orchestrating consecutive phases of cell growth (G1), DNA replication (S), DNA proofreading (G2), and cell division (mitosis, M) (Harper and Brooks, 2005). In mammalian cells, cell cycle progression is driven by cyclin-dependent kinases (CDKs) and their specific activators cyclins, while specialised molecular checkpoints ensure correct completion of each phase (Agarwal et al., 1995; Girard et al., 1991; Harper and Brooks, 2005; Lukas et al., 2004; Ohtsubo et al., 1995; Sherr, 1993, 1994; Sherr and Roberts, 1999, 2004; Vermeulen et al., 2003; Walker and Maller, 1991). If a problem occurs during replication, such as DNA damage, checkpoint activation serves to pause the cell cycle and repair the damage or, if not possible, to induce apoptosis (Agarwal et al., 1995; Awasthi et al., 2015; Lukas et al., 2004; Maréchal and Zou, 2013).

Given its central role In cell state, metabolic activity, availability of replication machinery and potential points of regulation, it is no surprise that viruses routinely target the host cell cycle to their own benefit (Fan et al., 2018). By dysregulating cell cycle checkpoints – to arrest or induce cell cycle progression – many viruses modify the cell environment to promote viral genome replication, protein production and the assembly of progeny virions (Bagga and Bouchard, 2014; Fan et al., 2018).

Vaccinia virus (VACV), the smallpox vaccine and prototypic poxvirus, is amongst the viruses shown to regulate the host cell cycle upon infection. Like all members of the *Poxviridae*, VACV is a large, double-stranded DNA virus which replicate exclusively in the cytoplasm of host cells (Moss, 2007; Condit et al., 2006; Dales and Siminovitch, 1961). To achieve this, VACV encodes and expresses a large subset of viral DNA replication proteins including a DNA polymerase, a helicase–primase, a uracil DNA glycosylase, a processivity factor, a single-stranded DNA-binding protein, a DNA ligase and the replicative protein kinase, B1 (Evans and Traktman, 1992; Moss, 2007).

Despite its exclusively cytoplasmic replication, VACV has been shown to inhibit cell cycle progression and mitosis (Jungwirth and Launer, 1968; Kit and Dubbs, 1962; Magee and Sagik, 1959; Magee et al., 1960). At a molecular level, infection was shown to alter expression of CDKs, cyclins, and the tumour suppressors Rb and p53 in pre-synchronized cells during late infection (Guerra et al., 2003; Wali and Strayer, 1999; Yoo et al., 2008). To date, the only VACV protein connected to cell cycle regulation is the viral kinase B1 (Santos et al., 2004). Overexpression of B1 in the absence of infection was found to mediate hyperphosphorylation of p53, thereby promoting its degradation. B1-mediated degradation of p53 was shown to require the E3 ligase Mdm2, which is the negative regulator of p53, and the proteasome (Haupt et al., 1997; Kubbutat et al., 1997; Santos et al., 2004). While B1-mediated degradation of p53 has not been investigated in the context of VACV infection, VACV was found to upregulate *Mdm2* transcription, suggesting that a similar p53 degradation mechanism employed during infection (Yoo et al., 2008).

As a crucial effector in checkpoint signalling, p53 can pause the cell cycle and induce apoptosis. Cellular stress, such as DNA damage, promotes phosphorylation and stabilisation of p53 which then directs transcription of target genes, including the cell cycle inhibitor p21 that can arrest cells in G1/S and G2/M (Agarwal et al., 1995; Dornan et al., 2003; Kastan et al., 1992; Sherr and Roberts, 1999; Taylor and Stark, 2001; Waldman et al., 1995). Conversely, inactivation of p53 prevents checkpoint-mediated cell cycle arrest in response to cellular stress, resulting in cell cycle dysregulation. Consistent with VACV-mediated dysregulation of the G1/S checkpoint through inhibition of p53 and Rb, a marked increase in S/G2 cells was observed during VACV infection (Yoo et al., 2008).

Despite downregulation of these checkpoint effectors, VACV was shown to activate the cellular DNA damage response (DDR) to facilitate host protein-assisted viral genome replication (Postigo et al., 2017). The cellular DNA single strand binding protein RPA was found to be recruited to replicating viral genomes where it acts as a platform for replisome assembly. Within the replisome, the viral polymerase E9 was shown to interact with the DDR-activating protein TOPBP1 and the cellular sliding clamp PCNA. While activation of the DDR kinases ATR and Chk1 was found to be essential, their role in VACV genome replication has not been established (Postigo et al., 2017). Additionally, mass spectrometry analysis of the proteins associated with replicating VACV genomes, suggested that VACV might exploit several other cellular DNA repair pathways including Non-Homologous End Joining, Base Excision Repair, Nucleotide Excision Repair, Interstrand Crosslink Repair, and Homologous Recombination Repair (HRR) (Reyes et al., 2017). While neither ATR nor Chk1 were detected, PCNA was strongly enriched, as well as all subunits of the mini-chromosome maintenance MCM2-7 replicative helicase complex, the HRR components BLM, MRE11, NBS1, and Ku70, as well as topoisomerase I/II which was previously shown to facilitate VACV replication (Lin et al., 2008).

Collectively these studies indicate that VACV infection triggers alterations in host cell cycle regulation and DDR to facilitate viral genome replication. However, the cellular factors and viral effectors that mediate these changes remain to be determined. By combining classical assays with state-of the art technologies we set out to investigate the effect of VACV on cell proliferation and characterise the (viral) effectors required to dysregulate the host cell cycle. Confirming earlier studies, we found that VACV both inhibits and shifts the host cell cycle (Jungwirth and Launer, 1968; Kit and Dubbs, 1962; Magee and Sagik, 1959; Wali and Strayer, 1999; Yoo et al., 2008). We further demonstrate that these two phenomena are independent of one another, with viral early genes responsible for inhibiting – and viral post-replicative gene(s) responsible for shifting – the cell cycle. Extending previous reports, we show that VACV B1/B12 mediates degradation of checkpoint effectors p53 and p21 to facilitate the cell cycle shift followed by DNA damage checkpoint activation by the viral kinase F10 (Postigo et al., 2017; Santos et al., 2004; Wali and Strayer, 1999; Yoo et al., 2008).

## RESULTS

### VACV infection inhibits cell proliferation

Cell proliferation is defined as the net change in cell number over time due to cell division and cell death. VACV infection has been reported to inhibit cell proliferation of mouse fibroblasts, and to have a pro-proliferative effect in human 143B osteosarcoma cells (Yoo et al., 2008). As this suggested that the effects of VACV infection on proliferation may be cell-type specific, we asked how VACV infection impacted proliferation of HeLa, BSC40 and HCT116 cells. Mock infected cells and cells infected with VACV were assessed at 0, 24 and 48 hours post infection (hpi) for cell numbers and cell death. While mock infected cell numbers doubled every 24h as expected, VACV infected cell numbers did not increase over the 48h time-course in any of the cell lines (Fig. 1A). The accompanying cytotoxicity assays showed that mock- and VACV-infected cells displayed the same level of cytotoxicity over 48h (Fig. 1B). These results indicated that VACV infection reduces cell proliferation through inhibition of cell division as opposed to induction of cell death.

**Figure 1.**
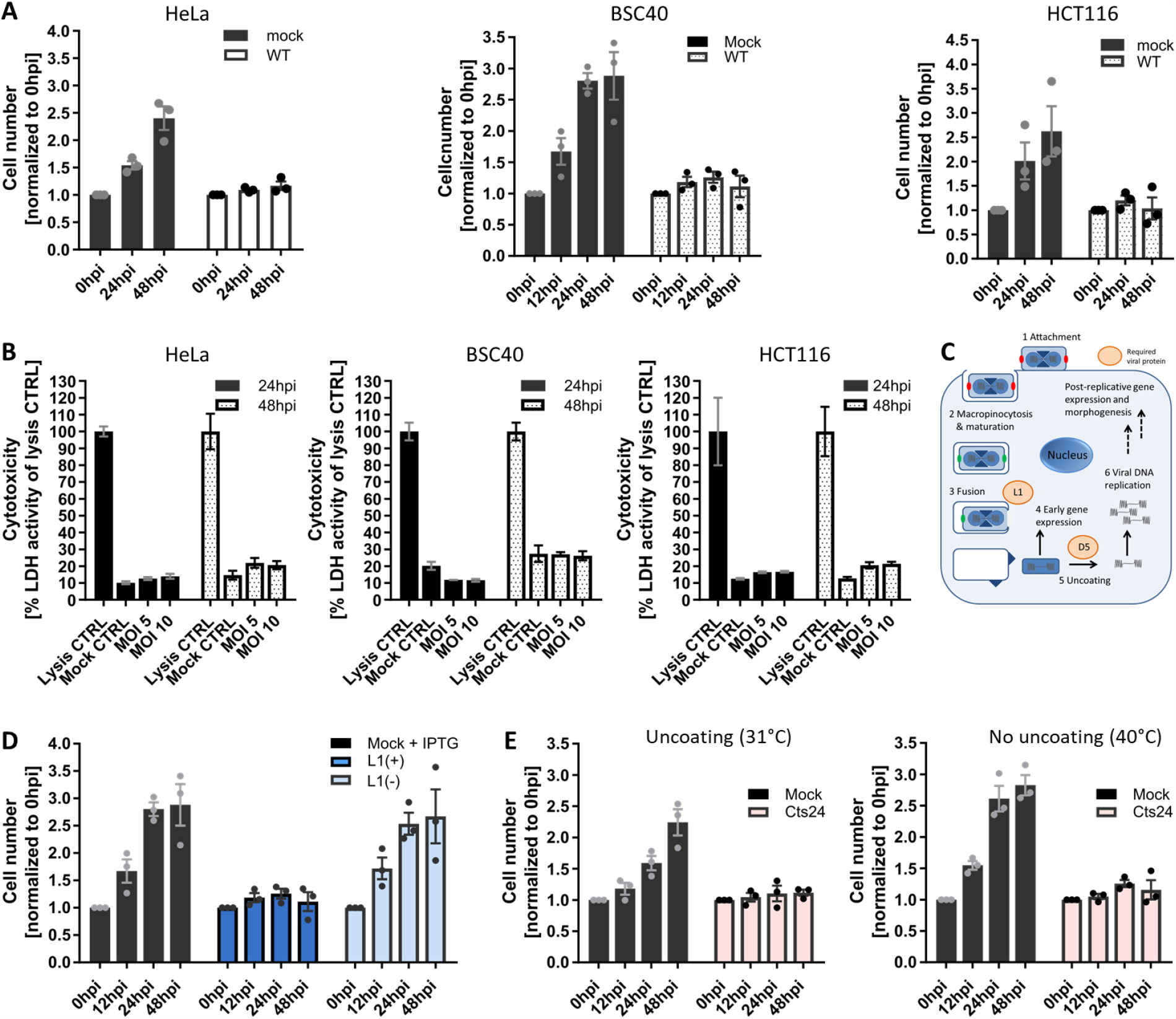
VACV inhibits the host cell cycle. [A] HeLa, BSC40, and HCT116 cells were either mock infected, or infected with WT VACV at MOI 5 and cell numbers were determined over 48 h. [B] HeLa, BSC40, and HCT116 cells were either mock infected, or infected with WT VACV at MOI 5, and MOI 10. Cytotoxicity was measured using the Pierce LDH Cytotoxicity Assay Kit at 24h and 48h and normalized to the cell lysis control. Data represent two biological replicates of technical triplicates and are displayed as mean ± S.D. [C] VACV life cycle schematic highlighting the viral proteins L1, and D5 which are required for fusion, and uncoating, respectively. [D] BSC40 cells were either mock infected, or infected with VACV L1(+), or VACV L1(-) at MOI 5 and cell numbers were determined over 48 h. [E] BSC40 cells were either mock infected, or infected with VACV C*ts*24 at MOI 5 and incubated at either the permissive (31°C), or non-permissive temperature (40°C) and cell numbers were determined during 48 h. [A, D, E] Data represent biological triplicates, each with technical duplicates, and is displayed as the normalized mean ± S.E.M.

### VACV-mediated inhibition of cell proliferation requires virus fusion but not genome uncoating

As illustrated in Figure 1C, VACV enters host cells by macropinocytosis (Chang et al., 2010; Mercer and Helenius, 2008). Once internalized, virions fuse from the macropinosome and deposit viral cores and their associated lateral bodies (LBs) into the host cytoplasm (Schmidt et al., 2013). Pre-replicative early gene expression is initiated allowing for the production of proteins required for subsequent genome uncoating (Kilcher et al., 2014), DNA replication and post-replicative intermediate and late gene expression (Moss, 2007). Using recombinant VACVs defective for virus entry or genome uncoating we sought to define the stage of the virus life cycle required for inhibition of cell division. To test whether VACV entry was required, we used a recombinant VACV (vL1i), which is inducible for the expression of the viral protein L1 (Bisht et al., 2008). The L1 protein is a member of the VACV entry-fusion complex, thus virions produced in the absence of L1, L1(-) virions, are unable to mediate membrane fusion (Bisht et al., 2008; Gray et al., 2019). When BSC40 cells were infected with L1(+) virions, cell proliferation was inhibited as expected (Fig. 1D). However, when cells were infected with L1(-) virions they continued to multiply at a level similar to mock infected cells, accumulating 2.7-fold over 48h. This result indicated that a post-fusion cytoplasmic stage of the VACV lifecycle is required to inhibit cell proliferation.

We next asked if inhibition of host cell proliferation occurred pre- or post-VACV genome uncoating using a VACV strain which is temperature sensitive (ts) for the expression of the viral uncoating factor D5 (C*ts*24) (Condit and Motyczka, 1981; Evans and Traktman, 1992; Kilcher et al., 2014). HeLa cells were infected with C*ts*24 under permissive (31°C) and non-permissive (40°C) uncoating conditions and cell proliferation was assessed over 48h. We found that VACV-mediated inhibition of cell proliferation was unabated under non-permissive uncoating conditions (Fig. 1E). This indicated that neither D5, nor the process of viral genome uncoating are required to block cell proliferation. Taken together, an event between VACV core deposition and genome uncoating is required to inhibit the host cell cycle.

### VACV infection induces a general cell cycle arrest

Having found that VACV inhibits cell proliferation we next asked at which cell cycle stage cells arrest upon infection. The cell cycle is divided into 4 consecutive phases: cell growth (G1), DNA replication (S), DNA repair (G2) and division (M). Cells that exit the cell cycle, enter quiescence (G0), a dormant state marked by the absence of cell proliferation, CDK activity and reduced levels of the DNA replication licensing factor MCM2 (Coller et al., 2006; Mlcochova et al., 2017; Musahl et al., 1998; So and Cheung, 2018; Terzi et al., 2016; Tsuruga et al., 1997).

Given the potent inhibition of cell proliferation observed upon VACV infection we first asked if VACV was driving cells into quiescence. We compared MCM2 protein levels between mock infected control and VACV infected cells over a 24h time course (Fig. 2A). We found that infection did not significantly alter MCM2 levels compared to mock infected cells (Fig. 2B). This suggested that VACV does not drive cells into quiescence but rather inhibits an active stage of the cell cycle.

**Figure 2.**
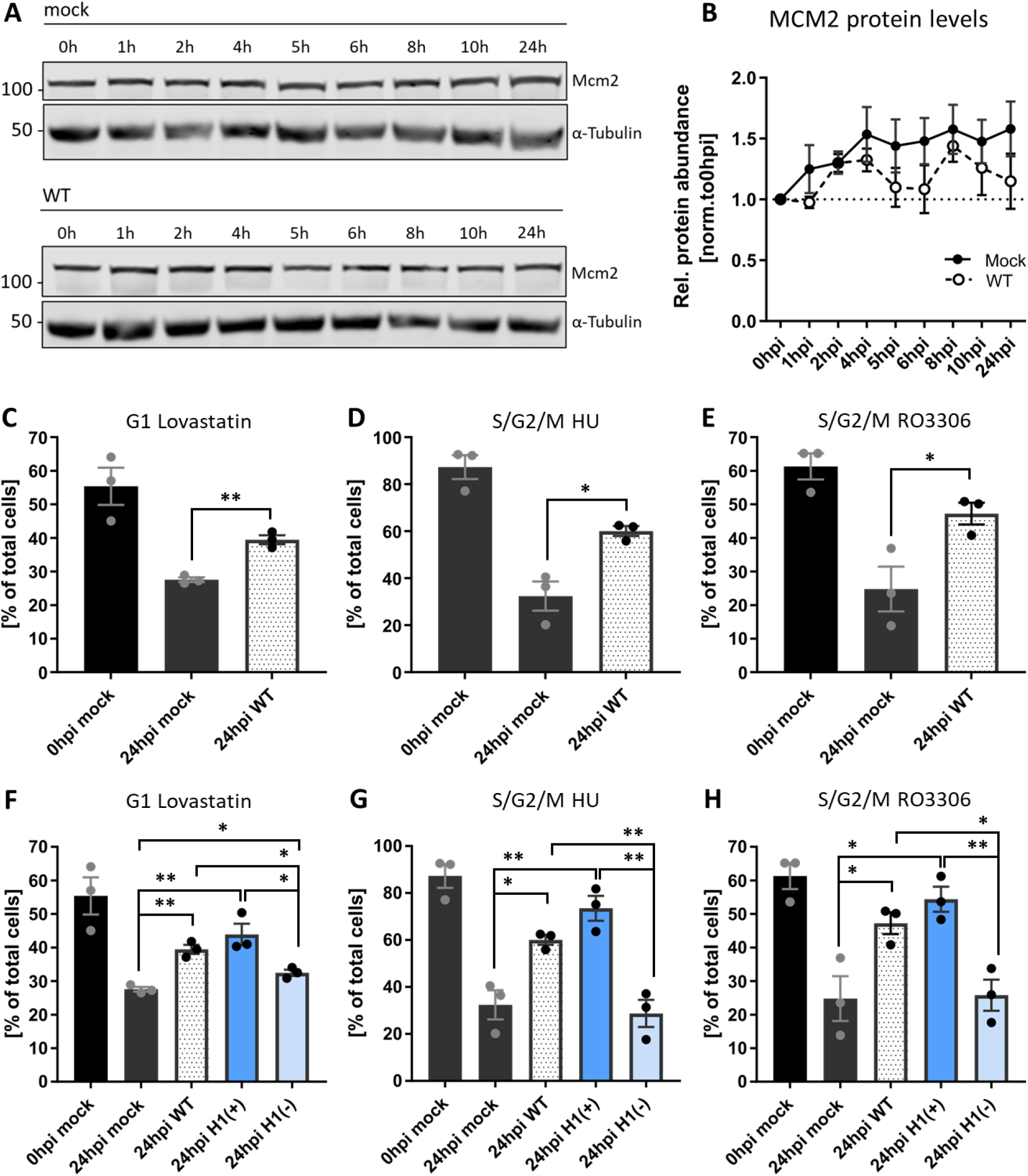
VACV establishes a general cycle block. [A] Immunoblot analysis of MCM2 during WT VACV infection. HeLa cells were either mock infected or infected with WT VACV at MOI 5 and harvested over 24h. Whole cell lysates were resolved via SDS-PAGE, and immunoblotted for MCM2, and α-Tubulin as loading control. A representative blot of three replicates is shown. [B] MCM2 protein abundance was quantified and normalized to the loading control. Data represents biological triplicates and is displayed as mean ± S.E.M. [C-H] HeLa FUCCI cells were synchronized in the indicated cell cycle stage with either Lovastatin, HU, or the CDK1 inhibitor RO3006. Immediately after release, cells were mock infected (black), or infected with WT (white), H1(+) (dark blue), or H1(-) (light blue) VACV at MOI 2. The cell cycle distribution was assessed at 0, and 24hpi by flow cytometry. [C, F] Percentage of G1 cells. [D-E, G-H] Percentage of S / G2 / M cells. All data represent biological triplicates and are displayed as mean ± S.E.M. Data in [C-E] is replicated in [F-H]. Parametric, unpaired, two-tailed t-test for significance. ns. p > 0.05, * p < 0.033, ** p < 0.0021.

To identify which stage(s) were inhibited, we used HeLa FUCCI cells which allowed us to distinguish four different cell cycle phases based on fluorescence: early G1 (no fluorescence), G1 (red only), early S (red and green), S / G2 / M (green only) (Fig. S1A,B) (Sakaue-Sawano et al., 2008). These cells were synchronized in either G1 with Lovastatin (Gray-Bablin et al., 1997; Rao et al., 1999), S phase with Hydroxyurea (Bacchetti and Whitmore, 1969; Fallon and Cox, 1979), or in late G2 with the CDK1 inhibitor RO3306 (Vassilev et al., 2006) (Fig. S1C-E). Cells were released, infected with VACV and the cell cycle distribution -relative to mock infected cells -determined at 24hpi. As expected, under all synchronization and release conditions (Lovastatin, Hydroxyurea and RO3306) mock infected cells re-entered the cell cycle resulting in reduced G1, S, and G2 fractions, respectively (Fig. 2**Error! Reference s ource not found**.C-E; black bars). Conversely, a portion of cells infected with VACV were retained in the pre-synchronized stage independent of the synchronizing agent: relative to mock infected cells 12% more VACV infected cells remained in G1 after release from Lovastatin (Fig. 2C), 28% more remained in S/G2/M after release from the Hydroxyurea S phase block (Fig. 2D) and 22% more remained in S/G2/M after release from the G2 block mediated by RO3306 (Fig. 2E). These results show that VACV inhibits G1, S, G2 and/or M phase progression suggesting that infection establishes a general cell cycle block.

As we found that a stage of the VACV lifecycle between fusion and uncoating inhibits cell proliferation (Fig. 1), and viral early gene expression (EGE) occurs between these two, we asked if EGE was required for the general block in cell cycle we observed. To address this, we used a recombinant VACV, v*ind*H1, which is inducible for the expression of the viral H1 phosphatase (Liu et al., 1995). Virions produced in the absence of inducer (H1(-)) are attenuated for early gene expression (Liu et al., 1995; Novy et al., 2018). As above, HeLa FUCCI cells were synchronised in G1, S or late G2 released, and infected with H1(+) or H1(-) virions. Cells were then assessed for their cell cycle distribution relative to WT infected cells. Cells infected with H1(+) virions, which are akin to WT virions, largely remained in their pre-synchronized cell cycle stage, while those infected with H1(-) virions were no longer capable of blocking cell cycle re-entry (Fig. 2F-H). Relative to H1(+) virions 11%, 45%, and 29% of cells re-entered the cell cycle after release from G1, S or late G2 blocks, respectively. This indicates that the H1 phosphatase, a known immune modulator (Najarro et al., 2001; Schmidt et al., 2013) -or viral EGE -is required for blocking cell cycle progression upon VACV infection.

### VACV infection inhibits cellular DNA synthesis

While we found that VACV can retain cells in G1 and S/G2/M, the FUCCI system did not allow for further resolution of the individual effect of VACV on S, G2 and M. Therefore, we next addressed whether VACV could inhibit M and/or S phase progression. To assess M phase during VACV infection, cells were infected with WT VACV, fixed, and stained with the DNA dye Hoechst at various time points between 0 and 24 hpi. Using genome condensation as a visual marker, we quantified the number of mitotic infected cells relative to the number of mitotic mock infected cells at each time point (Fig. 3A). While the percentage of mitotic cells fluctuated between 1.3% and 8.1%, the number of mitotic cells in infected samples declined until no mitotic cells were observed by 24hpi (Fig. 3A). This indicated that VACV is capable of blocking M phase either by preventing entry into mitosis, or by arresting cells prior to DNA condensation.

**Figure 3.**
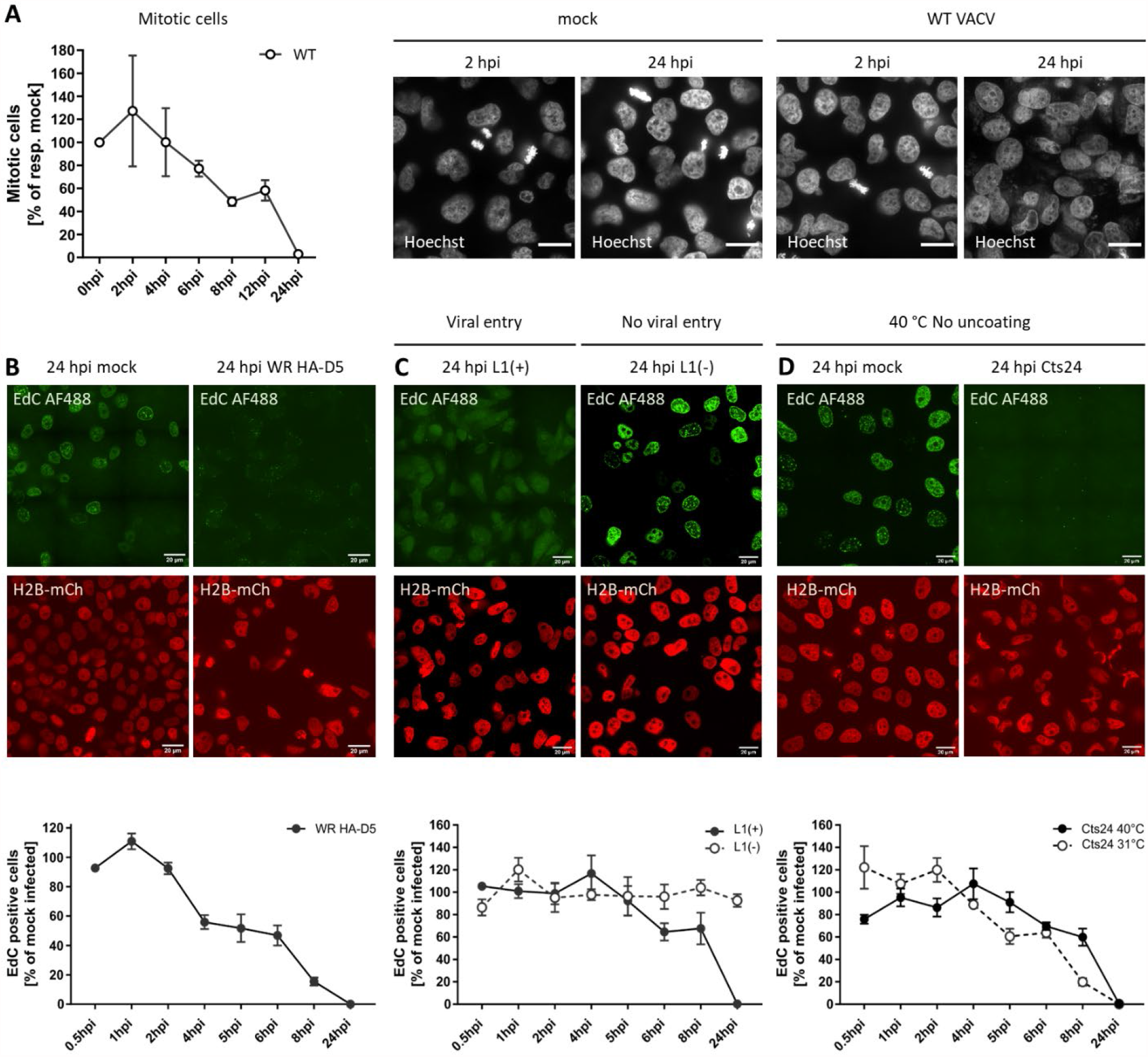
VACV inhibits mitosis and cellular DNA synthesis. [A] To quantify mitotic cells during VACV infection, HeLa cells were either mock infected, or infected with WT VACV at MOI 5, fixed and stained with the DNA dye Hoechst at various time points between 0 and 24 hpi. Using genome condensation as a visual marker, the number of mitotic infected cells was quantified relative to the number of mitotic mock infected cells at each time point. Representative images are shown. Scale bar = 20μm. Plotted data represent biological triplicates and are displayed as mean ± S.E.M. [B-D] Pulse-labelling with the nucleotide analogue EdC was used to identify active cellular DNA synthesis in VACV infected cells. HeLa H2B-mCh cells were mock infected, or infected with VACV at MOI 8 and pulse-labeled with the nucleotide analogue EdC (10 μM) for 15 min prior to fixation. Incorporated EdC was detected using the Click-iT^™^ AF488 Imaging Kit and imaged by confocal microscopy. Representative micrographs are shown. Scale bar = 20μm. Per condition and biological replicate, 150-450 nuclei were scored either as EdC positive, or negative and normalized to the respective mock infected control. Data represent biological triplicates and are displayed as mean ± S.E.M. [B] BSC40 cell were infected with WT HA-D5 VACV. [C] The requirement for viral entry was tested by infecting BSC40 cells with recombinant VACV L1(+) and L1(-) VACV. [D] To test if viral entry was required, BSC40 cells were infected VACV C*ts*24 and incubated at either the permissive (31°C), or non-permissive temperature (40°C).

We next addressed the impact of VACV infection on S phase, during which the cellular genome is replicated. It has been reported that VACV inhibits cellular DNA replication in HeLa cells (Jungwirth and Launer, 1968). To confirm and extend these findings, we established a microscopy-based cellular DNA replication assay using the nucleotide analogue EdC, which we have shown readily incorporates into cellular DNA, but not VACV replication sites (Wang et al., 2013). By pulse-labelling mock and VACV infected HeLa cells that express histone H2B-mCherry for 15 minutes prior to fixation and labelling, we could quantify the percentage of cells undergoing active cellular DNA synthesis (Illustrated in Fig. S2). When the percentage of EdC-positive infected cells was quantified over time we found that VACV infection completely blocked cellular DNA synthesis within 24hpi, consistent with previous reports (Fig. 3B). Inhibition of cellular DNA synthesis appeared to be due to an early event in the virus lifecycle with a 44% and 84% reduction in EdC positive infected cells, relative to controls at 4 and 8 hpi, respectively.

### Inhibition of cellular DNA synthesis is independent of viral genome replication

To determine which stage of the virus lifecycle was required for the observed inhibition of cellular DNA synthesis we repeated this assay with viruses defective for either virus fusion (vL1i) or genome uncoating (C*ts*24) as described in Fig. 1. We found that fusion competent L1(+) virions abolished EdC incorporation by 24 hpi, while fusion-incompetent L1(-) virions appeared to have no impact on cellular DNA synthesis relative to controls (Fig. 3C). As observed for inhibition of cell proliferation (Fig. 1), a post-fusion cytoplasmic stage of the virus lifecycle is also responsible for inhibition of cellular DNA synthesis.

Using C*ts*24, we next asked if genome uncoating was required to inhibit cellular DNA synthesis. Infection under both permissive uncoating (C*ts*24 31 °C) and non-permissive uncoating (C*ts*24 40 °C) conditions resulted in complete inhibition of EdC incorporation by 24hpi (Fig. 3D). This indicated that neither viral genome uncoating, nor viral genome replication are required to block cellular DNA synthesis. Collectively, these results suggest that viral core deposition and/or a lateral body constituent are required to block cellular replication.

### VACV post-replicative gene expression shifts the host cell cycle

Our results indicate that VACV blocks host cell proliferation, arresting cells in G1, S, and G2 and inhibits both mitosis and DNA replication (Figs. 1-3). This suggested to us that VACV either inhibits the cell cycle systemically (*i*.*e*., upon infection cells remain in the cell cycle stage they are in), or that VACV arrests DNA replication and mitosis, thereby effectively trapping cells in either G1 or G2 phases of the cell cycle. To test this, we infected unsynchronized HeLa FUCCI cells with WT VACV and followed the cell cycle distribution over 24h (Fig. 4A). Relative to mock infected cells, those infected with VACV gradually shifted from G1 into S/G2/M resulting in a significant increase in the S/G2/M fraction at the expense of G1 by 24hpi. Having established that infection inhibits cellular DNA replication (S phase) and mitosis (M phase), this result suggests that infected cells are shifted out of G1 and likely accumulate in S/G2.

**Figure 4.**
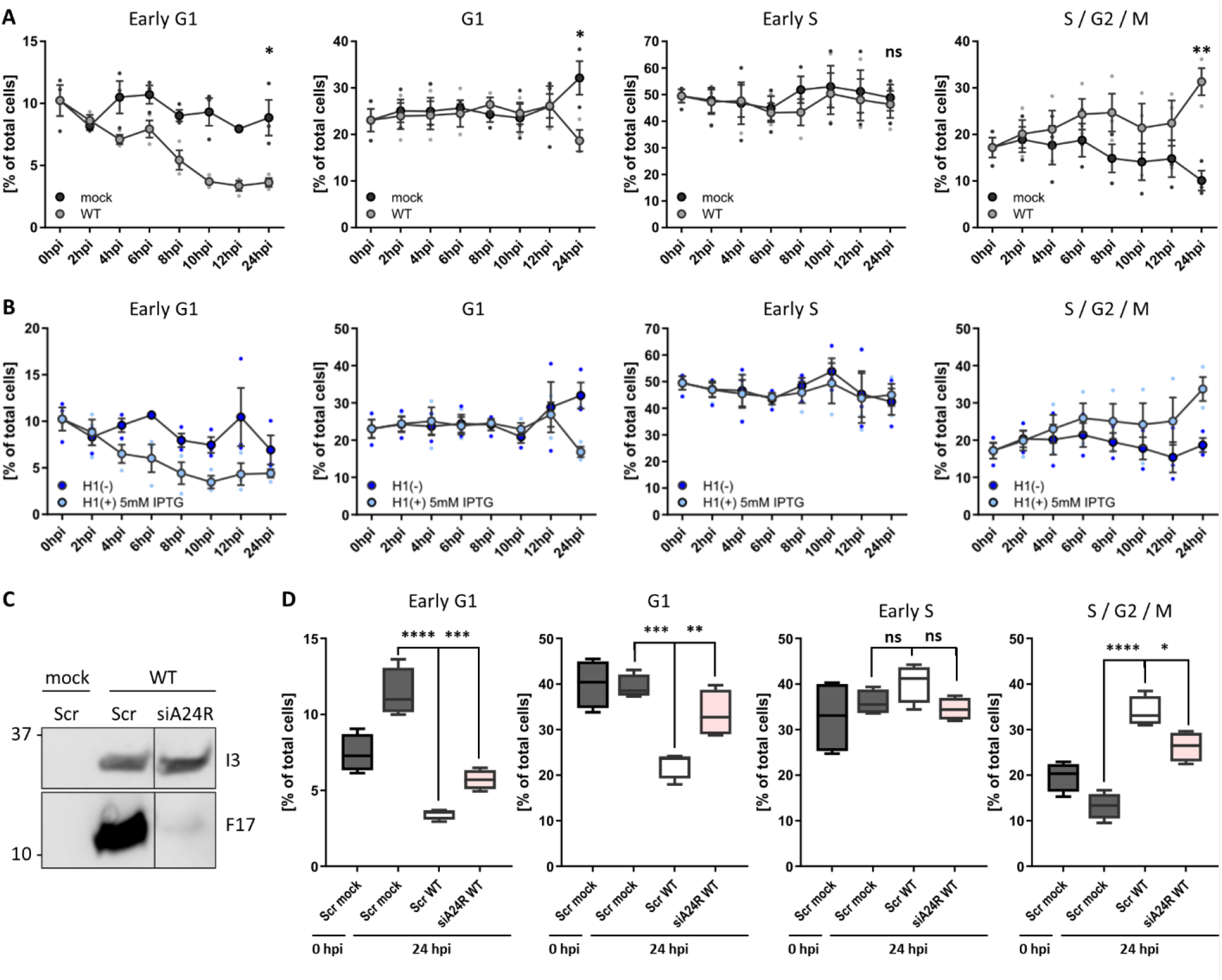
VACV post-replicative gene expression shifts the host cell cycle. [A] Cell cycle distribution of WT VACV infected HeLa cells compared to uninfected controls. HeLa FUCCI cells were either mock infected or infected with WT VACV at MOI 2 and samples were harvested between 0 h and 24 h. Cells were analysed by flow cytometry and the percentage G1, G1, early S, S / G2 / M were assessed. Data represent biological triplicates and are displayed as mean ± S.E.M. [B] Cell cycle distribution of VACV H1(-) infected HeLa cells compared to parental H1(+) infected controls. Samples were treated and analysed as in [A]. [C-D] To test if viral post-replicative gene expression was required to shift the cell cycle, the viral transcription factor A24 was silenced by siRNA. HeLa FUCCI cells were reverse transfected either with a scrambled control siRNA (Scr), or with siRNA targeting the viral transcription factor A24 (siA24R). Samples were infected with WT VACV at MOI 1 and harvested at 24 hpi. [C] Inhibition of viral post-replicative gene expression was validated by immunoblot analysis of the viral early gene I3, and the viral late gene F17. [D] The cell cycle distribution was determined by flow cytometry and compared to Scr mock (grey) and WT VACV (white) infected control samples. Percentage of HeLa FUCCI cell in early G1, G1, early S, and S / G2 / M. Data represent four biological replicates and are displayed as box (min to max), mean (line), and whiskers indicating the 1 to 99 percentiles. Parametric, unpaired, two-tailed t-test for significance. ns. p > 0.05, * p < 0.033, ** p < 0.0021, *** p < 0.0002, **** p < 0.0001

To define which step of the virus life cycle was required to shift the host cell cycle out of G1 we tested the requirement for viral EGE by following the cell cycle distribution of unsynchronized HeLa FUCCI cells infected with H1(+) or H1(-) VACV (Fig. 4B). As expected, H1(+) infected cells were gradually shifted out of G1 resulting in accumulation in S/G2 by 24hpi. In the absence of H1, VACV could no longer shift cells from G1 to S/G2 indicating that viral EGE and/or H1 was required to shift the host cell cycle. To narrow in, we asked if post-replicative viral gene expression could mediate the cell cycle shift. For this, we depleted a component of the VACV DNA-dependent RNA polymerase, A24, using virus targeting siRNA (Kilcher et al., 2014). As the VACV DNA-dependent RNA polymerase is pre-packaged in virions to direct in-core viral early gene transcription, siRNA depletion of A24 only affects post-replicative intermediate and late gene transcription which rely on newly expressed A24. The effectiveness of this approach was demonstrated by monitoring the expression of an early (I3) and late gene (F17) after A24 depletion (Fig. 4C). Relative to mock infected cells treated with control siRNA, WT VACV infection had shifted the distribution of cells from G1 into S/G2 by 24h as expected (Fig. 4D; Scr WT). Depletion of A24 prevented the shift from G1 to S/G2 in WT infected samples (Fig. 4D; siA24R WT). Compared to scrambled WT VACV controls, the G1 fraction was significantly increased, and the S/G2 fraction significantly decreased when viral post-replicative gene expression was inhibited. These findings suggest that viral post-replicative gene expression is required to shift the host cell cycle from G1 to S/G2 late in infection. Of note, the distinct viral requirements and timings suggest that the virus-induced cell cycle arrest described in figure 2 is independent from the cell cycle shift.

### The viral kinase F10 activates the cellular DDR late in infection

To gain additional insight into how VACV arrests and shifts the host cell cycle we turned to the underlying cell signalling pathways that control these processes. Cyclin-dependent kinases (CDKs) and their activators, the cyclins, are the key drivers of cell cycle progression. Periodic expression of cyclins targets the activity of their partner CDK to defined phases of the cell cycle. It has been shown that VACV alters CDK and cyclin expression in synchronized rabbit fibroblasts, as well as the transcript levels of cyclins in unsynchronised HeLa cells (Wali and Strayer, 1999; Yoo et al. 2008, Guerra et al. 2003). To test if VACV altered the expression of CDKs and Cyclins at the protein level in unsynchronised HeLa cells, we analysed CDK 1, 2, 4, 6, 7 and Cyclin A, B, D, E by immuno-blot analysis over 24h (Fig. S3). Contrary to previous findings with synchronised cells (Wali and Strayer, 1999), neither CDK (Fig. S3A) nor Cyclin (Fig. S3B) protein levels were found to be differentially regulated during the first 24h of infection (Fig. S3C).

As these results indicate that VACV does not prevent proliferation by depleting positive regulators, we next focused on cell cycle inhibitors. Molecular checkpoints can either pause the cell cycle or induce apoptosis upon aberrant cell cycle events such as DNA damage. The kinases ATR and/or ATM detect DNA breaks and trigger the cellular DNA damage response (DDR) through phosphorylation of the kinases Chk1 and/or Chk2 (Awasthi et al., 2015; Ciccia and Elledge, 2010; Maréchal and Zou, 2013). As VACV has been reported to activate the DDR in a pre-uncoating step (Postigo et al., 2017), we asked if DDR was the trigger for cell cycle arrest during early infection. First, we monitored DDR activation during VACV infection using phosphorylation of Chk1 (Ser345) and Chk2 (Thr68) as a readout. Immunoblot analysis of WT VACV infected HeLa cells showed Chk1 Ser345 phosphorylation after 8hpi, and Chk2 Thr68 phosphorylation from 4hpi, with a strong signal increase after 6hpi (Fig. 5A,B). DDR activation did not occur in mock controls (Fig. S4), and was not due to cellular DNA damage as determined by immunoblots directed against phosphorylation of γH2AX Ser139 (Fig. 5A). While these findings confirmed robust DDR activation in response to VACV infection, the kinetics of the response under our experimental setting was not consistent with a pre-uncoating event as the trigger (Postigo et al., 2017).

**Figure 5.**
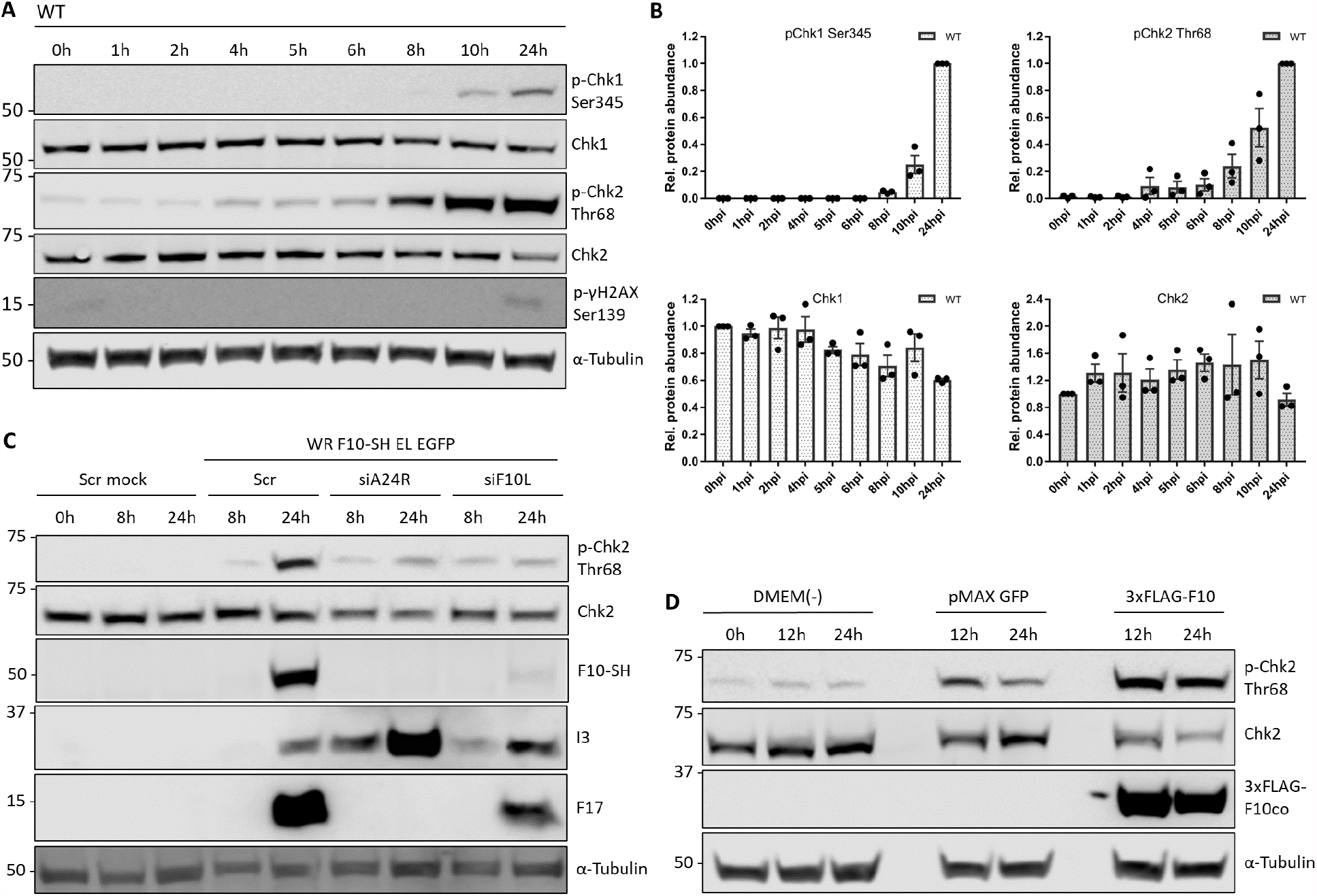
The viral kinase F10 activates the cellular DNA damage response. DDR activation was measured by phosphorylation of the DDR effector kinases Chk1 and Chk2, respectively. [A] HeLa cells were infected with WT VACV at MOI 5 and samples were harvested between 0 h and 24h. Whole cell lysates were resolved via SDS-PAGE and immunoblotted for activating phosphorylation of Chk1 (Ser345), Chk1, activating phosphorylation of Chk2 (Thr 68), phosphorylated γH2AX (Ser139) which marks cellular DNA damage, and α-Tubulin as loading control. A representative blot of biological triplicates is shown. [B] Total and phosphorylated protein abundance was quantified and normalized to the loading control. Data represent biological triplicates, normalized to 0 h, and 24h, respectively, and are displayed as mean ± S.E.M. [C] The late viral kinase F10 is required to activate the cellular DDR. HeLa cells were reverse transfected with scrambled control siRNA (Scr), siA24R, or siF10L. 48h after transfection, cells were either mock infected, or infected with WR F10-SH EL EGFP (MOI 1) and samples were harvested at 0, 8, and 24hpi. Whole cell lysates were resolved via SDS-PAGE and immunoblotted for pChk2 (Thr68), Chk2, HA to detect expression of viral F10-SH, the early viral protein I3, the late viral protein F17, and α-Tubulin as loading control. A representative blot is shown. [D] The late viral kinase F10 is sufficient to activate the cellular DDR. HeLa cells were transfected with either a DMEM(-) control, pMAX GFP control vector, or codon-optimized 3xFLAG-F10co. Samples were harvested at 0h, 12h, and 24h post transfection. Whole cell lysates were resolved via SDS-PAGE and immunoblotted for activating phosphorylation of Chk2 (Chk2 Thr68), Chk2, FLAG to monitor expression of 3xFLAG-F10co, and α-Tubulin as loading control. A representative blot of three biological replicates is shown.

The late phosphorylation of Chk1 and Chk2 suggested that a post-replicative event in the VACV life cycle activated the DDR. We therefore tested the requirement for viral intermediate and late gene expression by silencing A24 (siA24R). HeLa cells were infected with a recombinant VACV strain, WR F10-SH EL EGFP, that expresses a C-terminal streptavidin-HA tagged version of F10 from its endogenous locus and EGFP under a viral Early/Late promoter from the TK locus. While the scrambled control siRNA did not affect VACV mediated Chk2 activation, siA24R strongly reduced Chk2 Thr68 phosphorylation in infected cells (Fig. 5C). This implicated a viral intermediate/late protein in Chk2 activation. As activation is driven by phosphorylation, we hypothesized that the late VACV-encoded kinase, F10 (Lin and Broyles, 1994; Punjabi and Traktman, 2005), may be responsible. F10 was depleted using siRNA (siF10L) and the knockdown confirmed by immunoblot directed against the SH tag (Fig 5C; F10-SH). We found that depletion of F10 prevented Chk2 Thr68 phosphorylation suggesting that F10 is required for DDR activation.

To determine if F10 was sufficient to trigger Chk2 phosphorylation, or if it requires other viral factors, we expressed codon-optimized version of 3xFLAG tagged F10 (3xFLAGco) in uninfected cells and monitored Chk2 Thr68 phosphorylation. Expression of 3xFLAG-F10 resulted in a 2.4-fold increase in Chk2 Thr68 phosphorylation over the GFP vector control (Fig. 5D). These results show that VACV F10 can mediate phosphorylation and activation of the DDR kinase Chk2 in the absence of infection.

### VACV induces degradation of p53 and p21

Both DDR and the G1/S checkpoint pause the cell cycle through activation of effectors such as p53 and its transcriptional target, the CDK inhibitor p21 (Bunz et al., 1998; Dornan et al., 2003; Waldman et al., 1995). Under homeostatic conditions, the p53-specific E3 ligase, Mdm2, ubiquitinates p53 resulting in its degradation (Haupt et al., 1997; Kubbutat et al., 1997). In stressed cells, activation of p53 and accumulation of p21 arrests the cell cycle through CDK inhibition, and further prevents cellular DNA replication through inhibition of the DNA-polymerase processivity factor PCNA (Waga et al., 1994). Not surprisingly, modulation of p53/p21 activity and expression is a well-documented viral strategy to create a pro-viral environment (Bagga and Bouchard, 2014; Fan et al., 2018). VACV has been shown to reduce p53 levels during late infection in pre-synchronised cells (Wali and Strayer, 1999; Yoo et al., 2008).

We therefore asked if VACV infection altered Mdm2, p53 and p21 levels independent of DDR activation using immunoblot analysis of Mdm2, p53 and p21 in mock and WT infected cells over 24 h (Fig. 6A,B). In mock infected cells Mdm2, p53, and p21 protein levels fluctuated but were detectable during our experiment (Fig. S5A). Compared to mock infected cells, VACV caused a rapid reduction in Mdm2 proteins levels by 4hpi followed by downregulation of p53 and p21 by 6hpi (Fig. 6B and Fig. S5B). Of note, the downregulation of Mdm2, p53 and p21 all occurred prior to late viral gene expression (Fig. 6B; F17)

**Figure 6.**
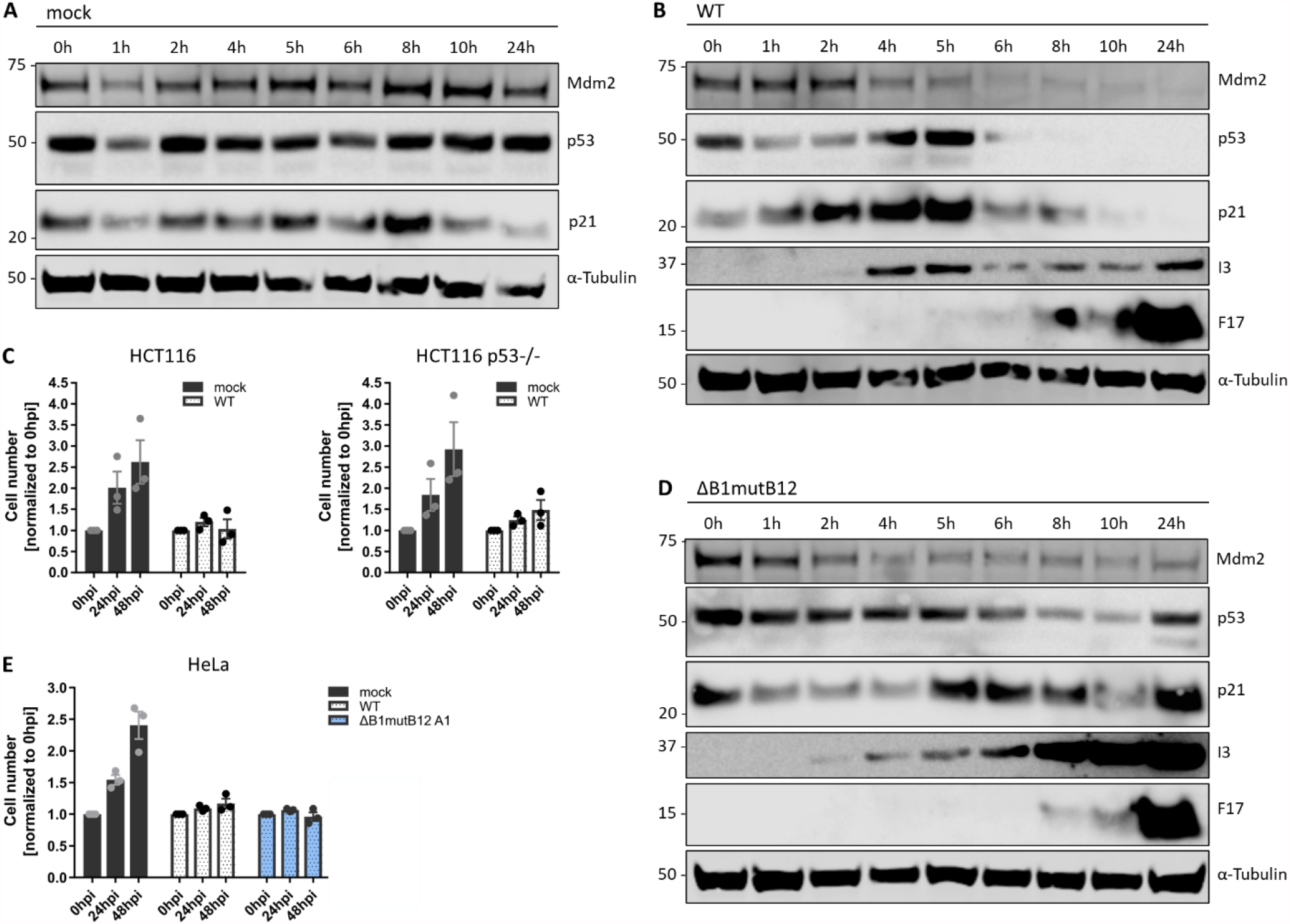
VACV infection degrades the DDR effectors Mdm2, p53 and p21. [A-B, D] HeLa cells were either mock infected, or infected with either WT VACV, or VACV ΔB1mutB12 at MOI 5 and harvested between 0 h and 24 h. Whole cell lysates were resolved via SDS-PAGE and blotted for Mdm2, p53, p21, the viral early protein I3, the viral late protein F17, and α-Tubulin as loading control. Representative immunoblots of three biological replicates are shown. [C] VACV inhibits cell proliferation independent of p53. WT HCT116 and HCT116 p53^-/-^ cells were either mock infected or infected with WT VACV at MOI 5 and cells were counted over 48 h. Cell numbers are normalised to 0 h and represent biological triplicates. Data is displayed as mean ± S.E.M. [E] VACV inhibits cell proliferation independent of B1 and B12. HeLa cells were either mock infected, or infected with either WT VACV, or VACV ΔB1mutB12 at MOI 5. Samples were processed and analyzed as in [C]. HeLa mock and WT data is reproduced from Figure 1 [A].

This finding implied that VACV does not arrest the cell cycle via upregulation of p53 and/or p21 expression. To exclude that p53 arrests the cell cycle prior to its downregulation at 6hpi, we asked if VACV could inhibit proliferation of HCT116 cells deleted of p53 (HCT116 p53-/-). HCT116 control and HCT116 p53-/-cells were mock infected or infected with WT VACV and cell numbers quantified at 24 and 48 hpi (Fig. 6C). As with BSC40 and HeLa cells, we observed a complete inhibition of cell proliferation upon infection of either cell line. These results confirmed that p53 is dispensable for VACV-induced cell cycle arrest.

### VACV ΔB1mutB12 fails to decrease cellular p53, and p21 levels

We next looked to identify the viral protein(s) that facilitate degradation of Mdm2, p53, and p21. In uninfected cells, phosphorylation of p53 at Ser15, Thr18, and Ser20 control its interaction with its negative regulator Mdm2 (Ashcroft et al., 1999; Hafner et al., 2019). As dynamic (de)phosphorylation regulates p53 stability, we hypothesized that a viral kinase or phosphatase may destabilize p53. The viral early kinase B1 was a promising candidate as p53 was degraded prior to late gene expression. In addition, B1 shares homology with the cellular kinase VRK1 (Vaccinia related kinase 1) which modulates p53 stability through Thr18 phosphorylation (Vega et al., 2004), and expression of B1 in uninfected cells was shown to be sufficient to phosphorylate p53 at its N-terminus, which was suggested to promote p52 binding to Mdm2 and proteasome-dependent degradation (Santos et al., 2004).

As B1 is essential for viral genome replication, siRNA silencing or genetic deletion (ΔB1) abrogate viral DNA synthesis and the production of VACV particles (Condit et al., 1983; Hooda-Dhingra et al., 1990; Olson et al., 2017). However, genetic truncation of the viral pseudokinase B12 in the ΔB1 virus (ΔB1mutB12) rescues viral replication through an unknown mechanism (Olson et al., 2019). Therefore, VACV ΔB1mutB12 allowed us to ask if B1 and/or B12 directed the downregulation of Mdm2, p53, and p21, independent of B1’s function in viral replication. Immunoblot analysis showed that, like WT VACV, ΔB1mutB12 caused a rapid decease in Mdm2 levels, however it failed to reduce the levels of p53, and p21 (Fig. 6D and S5C,D). This indicates that B1 and/or B12 mediate the degradation of p53, and p21 during VACV infection, whereas the reduction of Mdm2 is independent of these viral factors.

Next, we asked if B1/B12 were required to block the host cell cycle, despite their function in degrading p53 and p21. As before, HeLa cells were infected with WT or VACV ΔB1mutB12 and the cell number, relative to mock infected cells, quantified at 24 and 48hpi. ΔB1mutB12 blocked cell proliferation to the same level as WT VACV confirming that B1, B12 and down-regulation of p53 and p21 are -as expected-not necessary for the VACV-induced cell cycle arrest.

## DISCUSSION

In this study we investigated the impact of VACV infection on the host cell cycle in cancer cells. We show that VACV causes a general cell cycle arrest and shifts cells from G1 into S/G2. Our data suggests that these effects are two independent functions of viral early, and post-replicative gene expression, respectively. Looking for the underlying molecular mechanism, we found that the viral kinases B1 and F10 modulate the cell cycle checkpoint machinery. F10 was found to trigger the DNA damage checkpoint and B1/B12 caused degradation of the checkpoint effectors p53 and p21.

To characterise how VACV arrests the cell cycle, we used WT VACV and a set of viral mutants that block at defined stages of the virus lifecycle. Consistent with previous reports, we showed that VACV infection triggers a general cell cycle block between virus fusion and genome uncoating (Jungwirth and Launer, 1968). Using a virus that lacks H1 phosphatase we pin-pointed this block to either H1 within virions or early gene expression. It has been reported that UV-irradiated VACV cannot trigger cell cycle arrest (Tsung et al., 1996). Given that UV-irradiated virus only expresses a subset of early genes, and pre-packaged H1 should not be impacted by UV treatment, it is likely that an early gene(s) and not H1 mediate the cell cycle arrest.

We reasoned that the viral effector(s) could prevent cell cycle progression by inhibiting a positive cell cycle regulator, such as CDKs or cyclins, or conversely by activating a negative cell cycle regulator, such as a cell cycle checkpoint. We found that VACV infection had no impact on the protein levels of Cyclin A, B, D, E and CDKs 1, 2, 4, 6, 7 up to 24hpi, despite transcriptomic analysis showing decreased mRNA levels for Cyclins A2, C, D1, G1, and H1 at 16hpi in HeLa cells (Guerra et al., 2003). In general agreement with our data, proteomic analysis indicated that Cyclin A2, B1, B2, H, and CDKs 1, 2, 6, 7 levels were not altered by infection (Soday et al., 2019). Taken together, these data strongly suggest that VACV infection does not inhibit cell proliferation by actively diminishing Cyclin and/or CDK protein levels. This data also suggests that VACV-mediated cell cycle arrest is specifically driven by an early viral gene, and not due to a general effect, such as virus-mediated host shut-off (Parrish and Moss, 2007; Parrish et al., 2007; Pedley and Cooper, 1984).

Next, we addressed if VACV cell cycle inhibition was triggered by activation of a cell cycle checkpoint. Complementing the function of Cyclins and CDKs, these checkpoints can pause the cell cycle in case of stress signals such as cellular DNA damage. As VACV infection was reported to activate the cellular DNA damage response (DDR) before viral genome uncoating (Postigo et al., 2017), we tested if this caused the cell cycle arrest. Using Chk1 and Chk2 phosphorylation as a readout of DDR activation we found that DDR was triggered during late stages of infection (>10h), and that Chk1 and Chk2 phosphorylation was dependent upon the post-replicative viral kinase F10. As F10 is packaged into nascent virions it remains to be determined how F10 triggers DDR and if pre-packaged F10 is responsible for early VACV-induced cell cycle arrest.

To test if VACV infection triggered a DDR-independent checkpoint, we next focused on the effector p53 and its transcriptional target, the cell cycle inhibitor p21 (Agarwal et al., 1995; Dornan et al., 2003; Kastan et al., 1992; Sherr and Roberts, 1999; Taylor and Stark, 2001; Waldman et al., 1995). In line with previous studies, we found that VACV infection resulted in depletion of p53 and p21 in a B1/B12 dependent fashion (Olson et al., 2017; Santos et al., 2004; Wali and Strayer, 1999). Contrary to previous reports (Santos et al., 2004; Yoo et al., 2008), we found that Mdm2, a negative regulator of p53 stability, was also degraded. This suggest that VACV uses an alternative, Mdm2-independent strategy for p53/p21 degradation during infection. None-the-less using p53-/-HCT116 cells and a B1 deletion virus we conclusively show that neither B1, nor p53 is required for VACV-induced cell cycle arrest. As checkpoint activation can also induce a p53-independent arrest (Deeds et al., 2003; Park et al., 2017; Rao et al., 1999; Taylor and Stark, 2001), these findings do not exclude that VACV inhibits the cell cycle through triggering a checkpoint.

While p53 degradation does not explain the cell cycle arrest, it might facilitate the VACV-induced cell cycle shift. Consistent with previous observations in cells released from a G1 block, we demonstrate that VACV post-replicative gene expression shifted cells into S/G2 at the expense of G1 and M phase (Wali and Strayer, 1999; Yoo et al., 2008). For cells to be able to transition from G1 into S/G2, they need to be released from the initial cell cycle block, established during early infection. Our data is in line with a model in which VACV allows S phase entry by disassembling the G1/S checkpoint through deactivation of p53, p21 and Rb (Yoo et al., 2008). That p53 and p21 are degraded before post-replicative gene expression suggests that deactivation of the G1/S checkpoint is not sufficient to shift infected cells. In agreement with our observation that cell cycle activators (CDKs and cyclins) are not reduced, our model presumes that infected, arrested cells still have the ability to progress in cell cycle. Together, these findings indicate that viral early gene expression triggers a general cell cycle arrest, then specifically relieves the G1/S checkpoint before post-replicative gene expression actively promotes a G1 to S/G2 transition (Figure 7).

**Figure 7.**
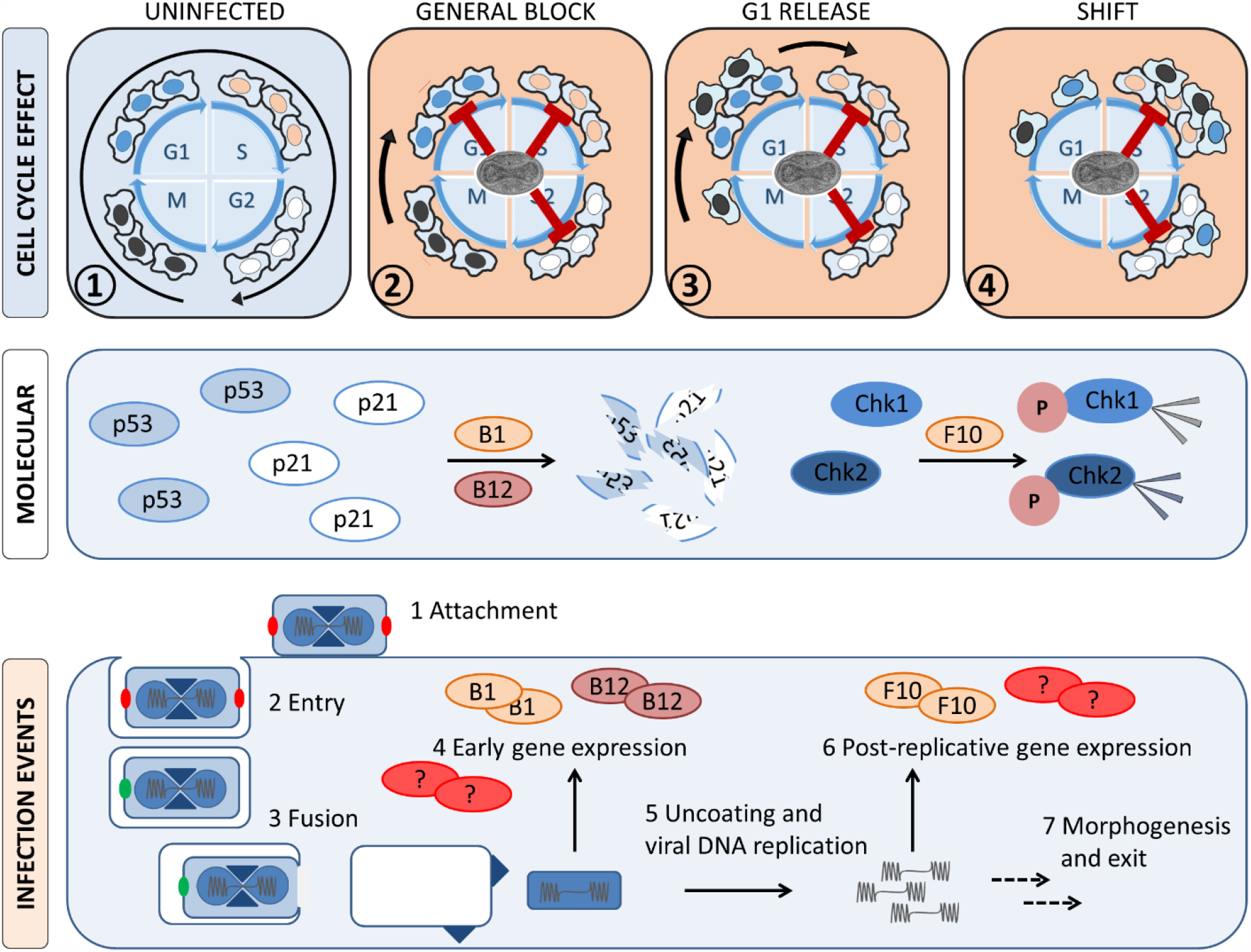
Model of VACV-induced cell cycle changes. VACV infection establishes a general cell cycle arrest early during infection. Progressing infection then selectively releases cells from the G1 block, which causes a cell cycle shift into S/G2. While the mechanism and cellular factors that regulate these changes are unknown, our data indicates that a viral early gene arrests the cell cycle and we show that the cell cycle shift is mediated by a post-replicative viral gene. We further demonstrate that the viral kinase F10 activates the cellular DNA damage response (DDR), while viral B1/B12 cause degradation of the DDR effector p53 and its transcriptional target p21. This observation is in line with the model that arrested, infected cells are released from the G1 block by inactivation of p53 and p21.

Taken together, we show that VACV inhibits the host cell cycle prior to genome uncoating, independent of the tumour suppressor p53, its transcriptional target p21 and DDR activation. We suggest that early degradation of p53 and p21 serve to deactivate the G1/S checkpoint allowing an unidentified viral late gene to shift the host cell cycle from G1 into S/G2. Future work will be aimed at identifying the viral “shift” protein, and determining if the initial cell cycle arrest is a direct effect of infection, or a more general effect of virus-induced host shutoff. Furthermore, the function of the observed cell cycle changes in the virus life cycle will be addressed.

## MATERIALS AND METHODS

### Cell culture

HeLa (human, ATCC), HeLa FUCCI (human, RIKEN Cell Bank, (Sakaue-Sawano et al., 2008)), HeLa H2B-mCh (human, kind gift from Dr. Serres), HCT116 (human, kind gift from Dr. Wilson), and HCT 116 p53^- /-^ (human, kind gift from Dr. Wilson) were grown in Dulbecco’s Modified Eagle Medium (DMEM, Life Technologies) supplied with 10% FBS (Life Technologies), 1% Gluta-Max (Gibco), 1% Penicillin-Streptomycin (Life Technologies), and 1% non-essential amino acids (Gibco). BSC40 (African green monkey) medium additionally contained 1% Sodium pyruvate (Life Technologies). Cells were maintained under 37.0 °C, 5% CO2.

### Antibodies and reagents

The following antibodies were used at 1:1,000 for western blotting, unless otherwise indicated; anti-CDK1 (Cell Signaling #9116), anti-CDK2 (Cell Signaling #2546), anti-CDK4 (Cell Signaling #12790), anti-CDK6 (Cell Signaling #13331), anti-CDK7 (Cell Signaling #2916), anti-Chk1 (Santa Cruz sc-56291), anti-phospho-Chk1 Ser345 (Cell Signaling #2348), anti-Chk2 (Cell Signaling #3440), anti-phospho-Chk2 Thr68 (Cell Signaling #2197), anti-CyclinA (Santa Cruz sc-71682), anti-CyclinB1 (Cell Signaling #4138), anti-CyclinD1 (Cell Signaling #2978), anti-CyclinE (BD Pharmingen 551160), anti-F17 (Liu et al., 1995), anti-FLAG (Abcam ab205606), anti-I3 (Rochester and Traktman, 1998), anti-HA (BioLegend 902302, 1:2,000), anti-HA (BioLegend 901502, 1:2,000), anti-phospho-histone γH2A.X Ser139 (Cell Signaling #9718), anti-MCM2 (Cell Signaling #12079), anti-Mdm2 (Abcam ab16895), anti-mouse-HRP (Cell Signaling #7076, 1:5,000), anti-mouse-IRDye 680RD (LI-COR 926-68072, 1:10,000), anti-mouse-IRDye 800CW (LI-COR 926-32212, 1:10,000), anti-p21 (Cell Signaling #2947), anti-p53 (Abcam ab1101), anti-rabbit-HRP (Cell Signaling #7074, 1:5,000), anti-rabbit-IRDye 680RD (LI-COR 926-68073, 1:10,000), anti-rabbit-IRDye 800CW (LI-COR 926-32213, 1:10,000), anti-α-Tubulin (Cell Signaling #3873, 1:2,000), and anti-α-Tubulin (Cell Signaling #2125, 1:2,000).

Inhibitors were used at the following concentrations: Mevinolin (Lovastatin, 20 µM, LKT labs), DL-Mevalonic acid lactone (Mevanolate, 6 mM, Sigma-Aldrich), Hydroxyurea (HU, 2.5 mM, Sigma-Aldrich), RO3306 (10 µM, Sigma-Aldrich). Mevinolin (Lovastatin) was converted to the active compound by dissolving the prodrug in 70% EtOH (JavanMoghadam-Kamrani and Keyomarsi, 2008). Additionally, Isopropyl β-D-1-thiogalactopyranoside (IPTG) was purchased from Sigma-Aldrich and used at the specified concentrations.

### Viruses, VACV purification, titration and infections

All viruses in this study were derived from the VACV Western Reserve (WR) strain. Apart from the wild-type (WT WR), we used recombinant WR HA-D5 (Kilcher et al., 2014), WR F10-SH EL EGFP, and WR ΔB1mutB12 (a kind gift of Dr. Wiebe (Olson et al., 2019)); temperature-sensitive WR C*ts*24 (a kind gift of Dr. Condit (Condit et al., 1983; Kilcher et al., 2014)); IPTG-inducible v*ind*H1 (a kind gift of Prof. Traktman, (Liu et al., 1995)), and vL1Ri EL EGFP (a kind gift of Dr. Moss, (Bisht et al., 2008)). WT and recombinant viruses were produced in BSC40 cells, and purified as previously described (Mercer and Helenius, 2008). The same protocol was followed for WR C*ts*24 but at permissive 31.0 °C. IPTG-inducible viruses were produced +/-IPTG (v*ind*H1: 5 mM, vL1Ri EL EGFP: 50 µM).

MV stocks were titered by plaque assay. Briefly, confluent BSC40 cells were infected with 10-fold serial dilutions of purified virus. After 48h, cells were fixed and stained with 0.1% crystal violet in 3.7% PFA. Temperature-sensitive WR C*ts*24 was titered under permissive (31.0°C) and non-permissive conditions (39.7°C). IPTG-inducible viruses were titered in medium supplied with IPTG. As v*ind*H1 and vL1Ri produced without IPTG (H1(-) and L1(-)) cannot be titered by plaque assay, virion concentration was determined indirectly through the DNA absorbance at 260/280nm of the virus stock (NanoDrop, Thermo Scientific). The MOI equivalent (MOI eq.) was then calculated using the 260/280 nm absorbance of WT VACV with a known pfu/ml. Cells were infected with the specified viruses and MOIs in DMEM without FCS (Life Technologies) for 1h at 37.0°C (C*ts*24: 31.0°C, and 39.7 °C). The inoculum was then replaced with supplemented medium and cells were incubated as indicated. For H1(+) and L1(+), both the infection and growth medium were supplied with the indicated concentration of IPTG.

### Cell proliferation assay

Cells were counted using the automatic cell counter Cellometer (Nexcelom) and seeded into 6-well plates (Greiner Bio-one). After incubation overnight samples were either mock infected or infected with the specified virus (MOI 5). Cells were harvested and cell counts were determined by two individual measurements per sample using the Cellometer (Nexcelom). To assess relative cell proliferation, cell counts were normalized to the respective baseline at 0hpi.

### Cytotoxicity assay

Cytotoxicity was determined with the Lactate Dehydrogenase assay (Pierce^™^ LDH Cytotoxicity Assay Kit, Thermo Scientific) as per manufacturer’s instructions. Cells grown to confluency in in 96-well plates were either mock infected or infected with the indicated virus (MOI 5 and MOI 10). At the indicated time post infection, the LDH assay was performed and absorbance measured at 490 nm and 650 nm with a VersaMax Microplate Reader (Molecular Devices) using SoftMax Pro software. The 490nm absorbance signal was background corrected by subtracting the absorbance at 650nm. Values were normalised against the lysis control.

### Western blotting and quantification

For immunoblots, cells were collected by scraping and centrifugation followed by resuspension in lysis buffer with protease inhibitors. Cell lysis was performed for 30min on ice, samples were centrifuged at 20,000g, 4 °C, 10min and the supernatant was boiled for 5min in 4x loading dye with 4x DTT prior to loading on 4-12% Bis-Tris polyacrylamide gels (Thermo Fisher Scientific). After transfer to nitrocellulose, membranes were blocked with 5% milk in TBS-T (Sigma Aldrich) for 1h before blotting. Primary antibodies were diluted in blocking solution and incubated overnight at 4 °C. Membranes were washed and incubated with HRP, or IRDye-secondary antibodies in blocking solution for 2h at RT, washed and analysed with ImageQuant LAS 4000 Mini (GE Life Sciences) and Luminata Forte Western HRP Substrate (Merck) for detection. IRDye secondary antibodies were imaged with a LiCOR ImageQuant. Independent of the detection method, protein intensities were quantified using the software StudioLite (LiCOR). Where applicable, samples were normalized against an internal α-Tubulin loading control.

### Cell cycle analysis by flow cytometry

Asynchronous, subconfluent HeLa FUCCI cells were either mock infected, or infected with the specified virus (MOI 2). Cells were washed, harvested and analysed using a GUAVA easyCyte HT (Merck Millipore, UK). Data was processed with InCyte 3.1.1 (Merck Millipore, UK), scoring cells as either early G1, late G1, early S, or S/G2/M (Fig. S1B, (Sakaue-Sawano et al., 2008)).

For synchronisation/release experiments, HeLa FUCCI cells were either arrested in G1 with Lovastatin (20µM), or in S phase with HU (2.5 mM), or in G2 with RO3006 (10 μM), as previously described (JavanMoghadam-Kamrani and Keyomarsi, 2008; Ma and Poon, 2011). Immediately before infection cells were released from the block by washing twice with PBS. For Lovastatin, release media was supplemented with Mevanolate (6mM). The samples were then prepared for flow cytometry and analysed as above.

### Visualisation of cellular DNA synthesis assay

HeLa Kyoto H2B-mCh cells seeded on coverslips were either mock infected or infected with WR HA-D5 (MOI 8). 15 min before fixation, cells were fed with 200 μl 4x EdC (40 μM, final = 10 μM). Cells were washed with PBS and fixed with 4 % PFA at RT for 15min. Fixed samples were washed 3x with PBS and stained for EdC using the Click-iT^™^ EdU Alexa FluorTM 488 Imaging Kit (Invitrogen), following manufacturer’s instructions. Coverslips were mounted with Immu-Mount (Thermo Scientific), and a minimum of 5 different locations per coverslip were imaged with a 100x oil immersion objective (NA 1.45) on a VT-iSIM microscope (Visitech; Nikon Eclipse TI). Nuclei were counted manually using ImageJ (minimum 450 nuclei per sample and biological repeat).

### Overexpression of viral proteins

Cells were transfected with 2µg of indicated plasmid DNA using Lipofectamine 2000 (Invitrogen) following manufacturer’s instructions.

### siRNA knockdown of viral proteins

Cells were reverse-transfected with the indicated siRNAs using Lipofectamine RNAiMAX (Invitrogen) following manufacturer’s instructions. Briefly, siRNA (stock 10 µM) and RNAiMAX were individually diluted in DMEM(-), combined and incubated for 1hr at RT. 250,000 cells in suspension were added to the siRNA / RNAiMAX solution (final siRNA concentration of 20 nM) and plated. Transfected cells were grown for 48h before infection. The following siRNA sequences with a dTdT modification (Sigma Aldrich) were used: siA24R 5’-CUGCUAAGCCGUACAACAA-3’ (Kilcher et al., 2014), siF10L 5’-AACUGGUAUUACGAUUUCCAUU-3’, Scrambled (allStar Negative).

## ACKNOWLEDGEMENTS

J. Mercer is supported by core funding to MRC Laboratory for Molecular Cell Biology at University College London (MC_UU12018/7) and the European Research Council (649101-UbiProPox). C. K. Martin is funded by the MRC Laboratory for Molecular Cell Biology PhD program. We thank Janos Kriston-Vizi for imaging assistance (MRC-UCL University Unit Grant Ref MC_U12266B). We thank Dr. Murielle Serres, Dr. Miranda Wilson and Dr. Matt Wiebe for reagents.

The authors declare no competing financial interests.

Author contribution: C. K. Martin and J. Mercer designed the study. C. K. Martin performed and analysed the experiments. J. R. Samolej performed the cytotoxicity assays. C. Bertoli, M. Wiebe, and R. deBruin provided key reagents and valuable comments throughout the project. C.K. Martin and J. Mercer wrote the manuscript with input from the other co-authors.

## Supplementary Figure legends

**Figure S1.**
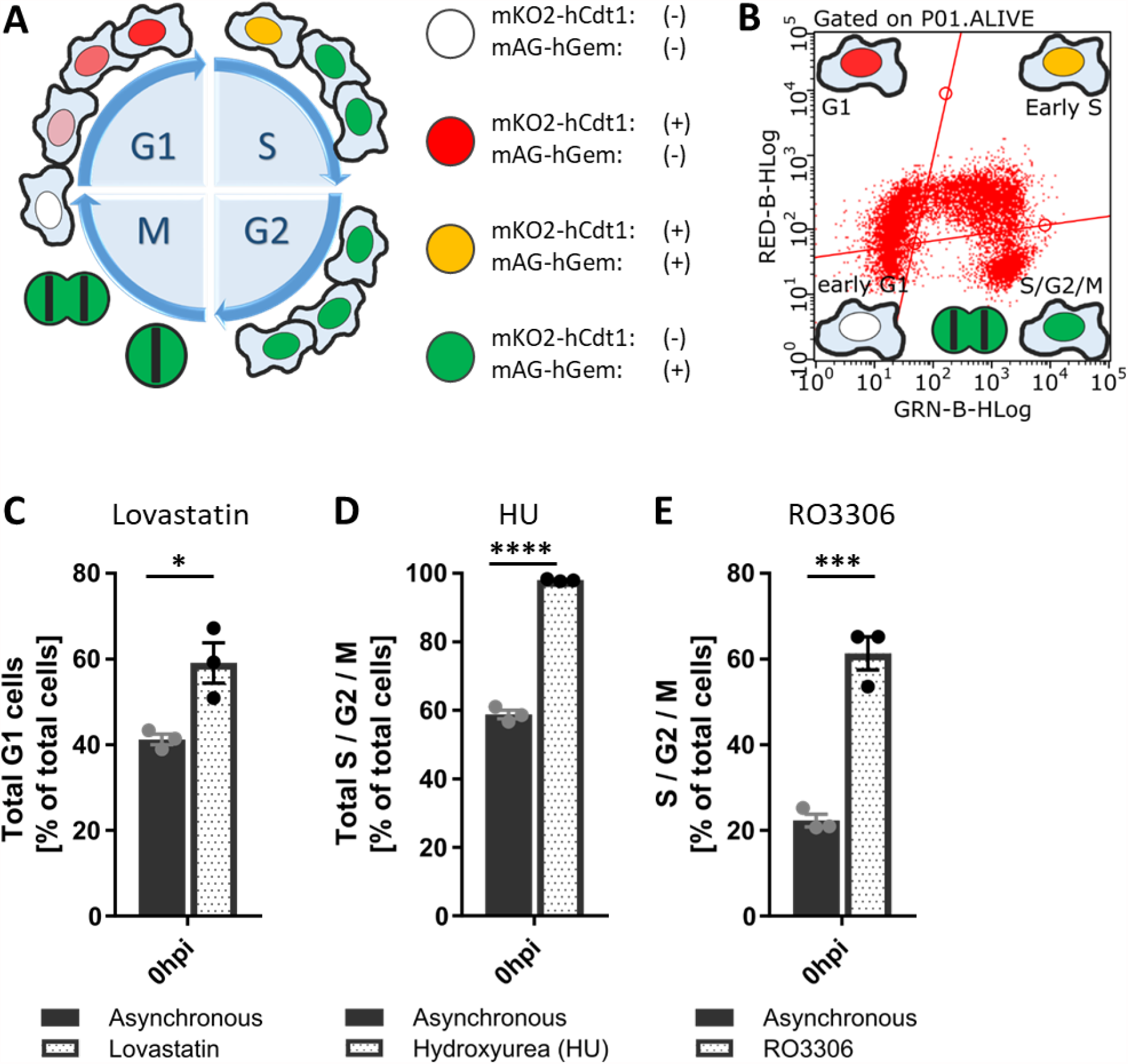
The HeLa FUCCI system for cell cycle analysis of synchronized and unsynchronized cells. [A] The HeLa FUCCI cell line stably expresses fluorescently tagged fragments of human Geminin (mAG-hGem(1-100), green) and Cdt1 (mKO2-hCdt1(30-120), red). Due to the antiphasic expression of the fragments, the relative abundance marks distinct phases of the cell cycle. The schematic shows cell cycle stage-dependent changes in fluorescence of the HeLa FUCCI cell line. Adapted from (Sakaue-Sawano et al., 2008). [B] Schematic flow cytometry fluorescence plot indicating the gating strategy used to group cells as either early G1 (no fluorescence), G1 (red), early S (green + red = yellow), and S/G2/M (green). [C-E] Synchronization effectiveness of Lovastatin, HU, and RO3306. [C] HeLa FUCCI cells were synchronized in G1 with Lovastatin (20 μM) for 24 h. Using flow cytometry, cells were classified as early G1, G1, early S, or S / G2 / M. Data represent the total G1 fraction (early G1 + G1) of either asynchronous, or Lovastatin treated cell populations. [D] HeLa FUCCI cells were synchronized in S phase with HU (2.5 mM) for 16 h and samples were analysed as in [C]. Data represent the total S / G2 / M fraction of either asynchronous, or HU treated cell populations. [D] HeLa FUCCI cells were synchronized in G2 phase with the CDK1 inhibitor RO3306 (10 μM) for 16 h and samples were analysed as in [C]. Data represent the S / G2 / M fraction (not including early S) of either asynchronous, or RO3306 treated cell populations. Experiments were conducted in biological triplicates and are displayed as mean ± S.E.M. Parametric, unpaired, two-tailed t-test for significance. ns. p > 0.05, * p < 0.033, ** p < 0.0021, *** p < 0.0002, **** p < 0.0001.

**Figure S2.**
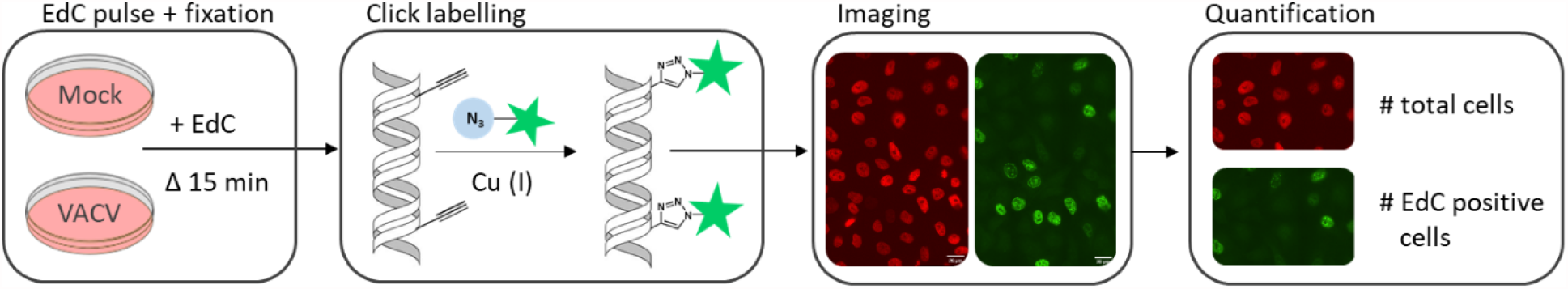
Experimental strategy to visualise cellular DNA synthesis in VACV infected cells. HeLa H2B-mCh cells were either mock infected or infected with VACV at MOI 8. 15 min before fixation, cells were pulse-labelled with the nucleotide analogue EdC (10 µM final). To detect nuclei with active DNA synthesis, incorporated EdC was covalently linked to AlexaFluor488 using the Click-iT^™^ Kit. Samples were imaged by confocal microscopy and the total amount of nuclei, marked by mCherry expression, as well as the number of EdC positive nuclei, marked by AF488 fluorescence, was determined.

**Figure S3.**
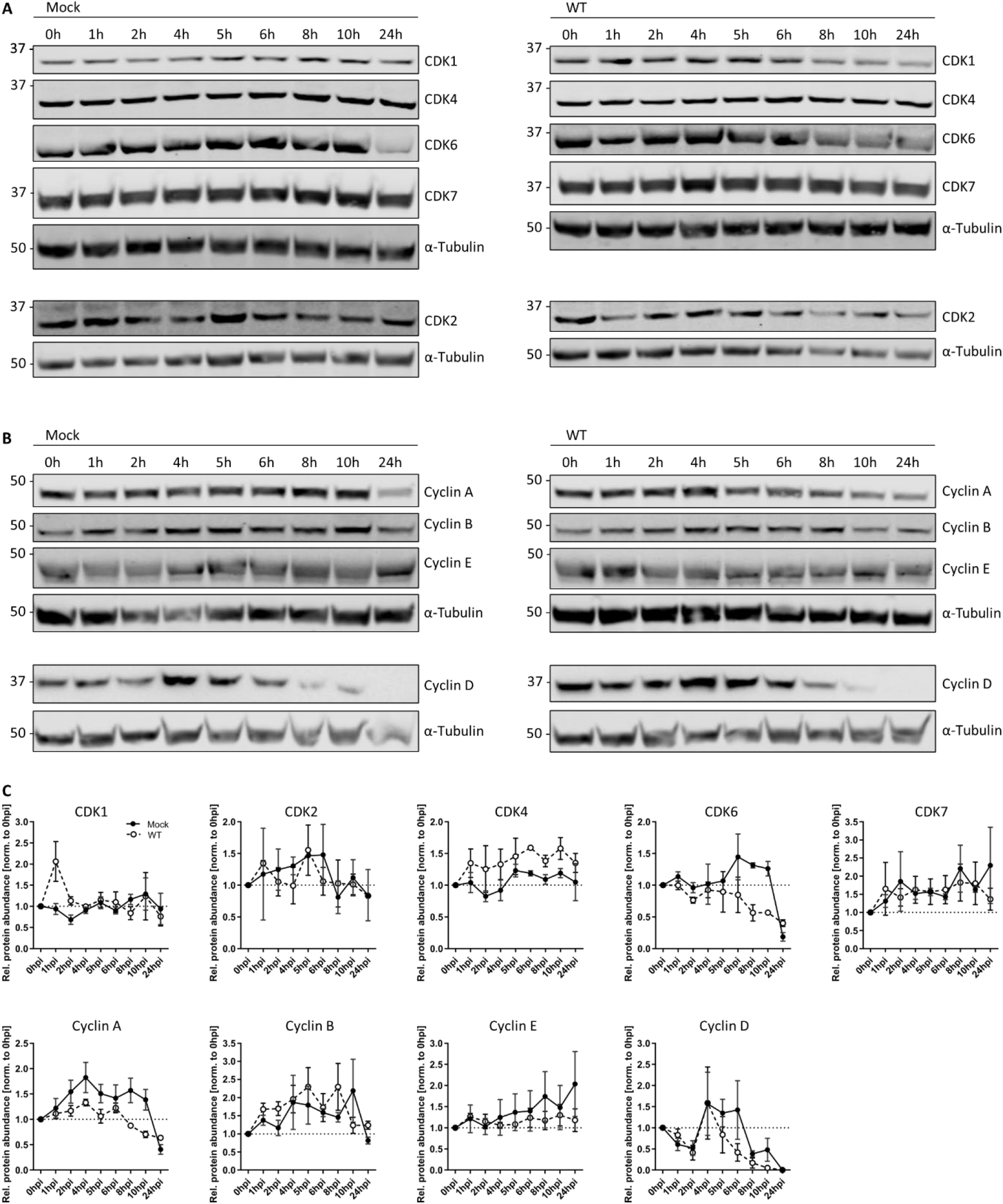
CDK and cyclin protein levels during VACV infection. [A-C] HeLa cells were either mock infected or infected with either WT VACV at MOI 5 and harvested between 0 h and 24 h. Whole cell lysates were resolved via SDS-PAGE and blotted for CDK 1, 2, 4, 6, 7, cyclin A, B, D, E, and α-Tubulin as loading control. [A-B] A representative blot of three biological replicates is shown. [C] Protein abundance was quantified and normalized to the loading control. Data represent biological triplicates, normalized to 0 h, and are displayed as mean ± S.E.M.

**Figure S4.**
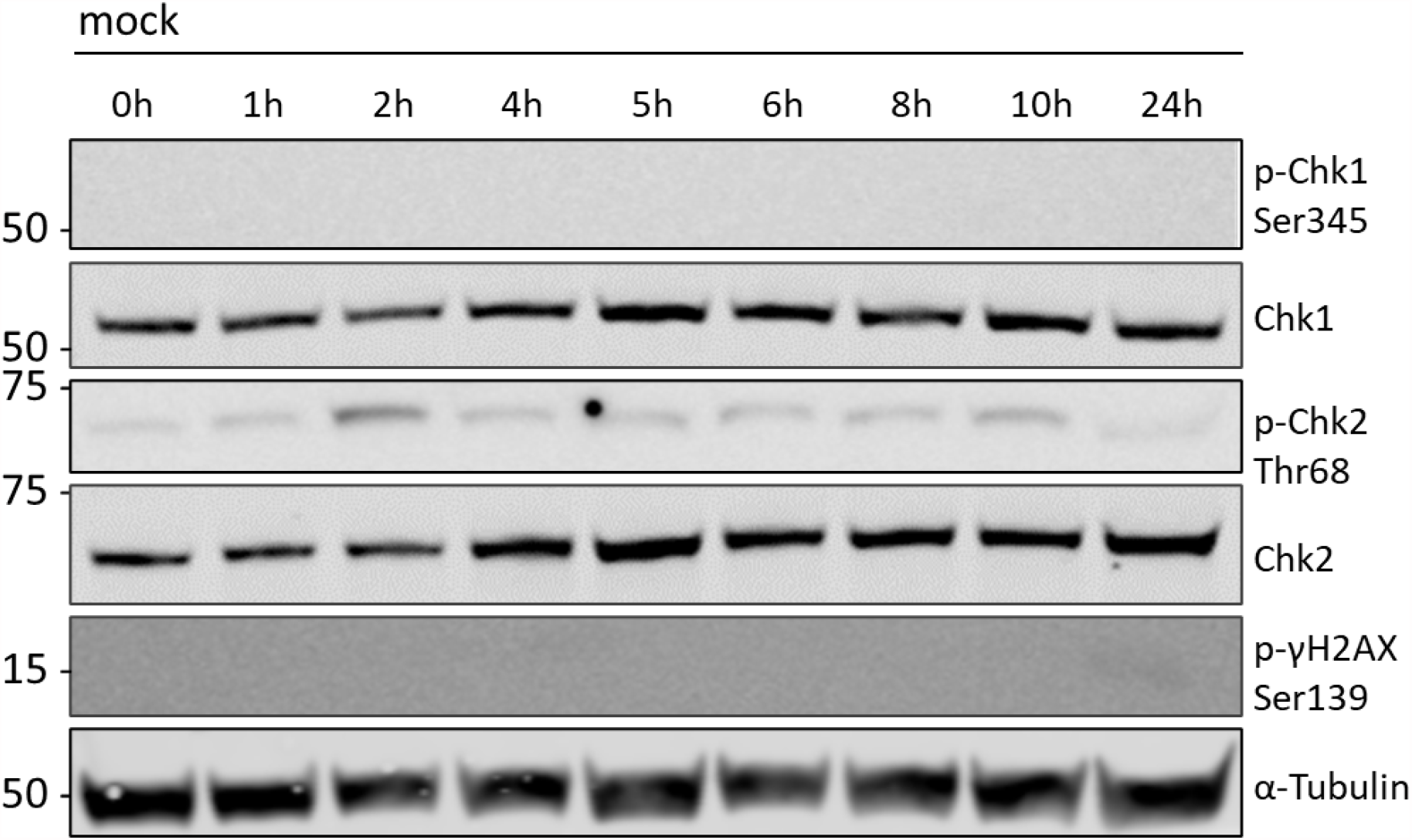
DDR activation level in mock infected HeLa cells. DDR activation was measured by phosphorylation of the DDR effector kinases Chk1 and Chk2, respectively. HeLa cells were mock infected and samples were harvested between 0 h and 24h. Whole cell lysates were resolved via SDS-PAGE and immunoblotted for activating phosphorylation of Chk1 (Ser345), Chk1, activating phosphorylation of Chk2 (Thr 68), phosphorylated γH2AX (Ser139) which marks cellular DNA damage, and α-Tubulin as loading control. A representative blot of biological triplicates is shown.

**Figure S5.**
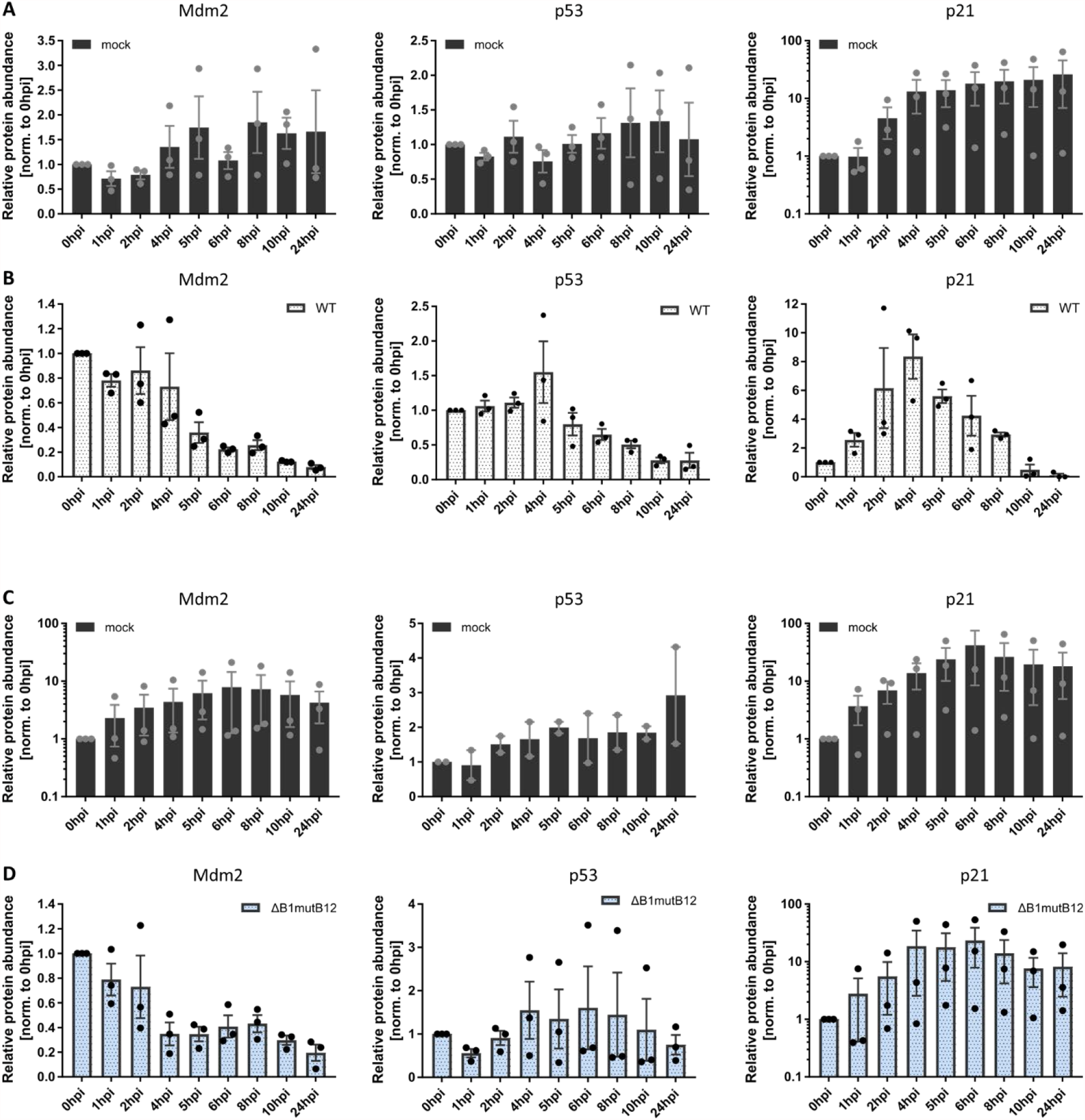
Quantification of Mdm2, p53, and p21 protein levels. [A-D] HeLa cells were either mock infected, or infected with either WT VACV, or VACV ΔB1mutB12 at MOI 5 and harvested between 0 h and 24 h. Whole cell lysates were resolved via SDS-PAGE and blotted for Mdm2, p53, p21, and α-Tubulin as loading control. Protein abundance was quantified and normalized to the loading control. Data represent biological triplicates, normalized to 0 h, and are displayed as mean ± S.E.M. [A] Mock infected control for WT VACV infection [B]. [C] Mock infected control for VACV ΔB1mutB12 infection [D].

## Notes

### Competing Interest Statement

The authors have declared no competing interest.

